# A human induced pluripotent stem cell toolbox for studying sex chromosome effects

**DOI:** 10.1101/2025.02.28.640277

**Authors:** Ruta Meleckyte, Wazeer Varsally, Jasmin Zohren, Jerry Eriksson, Tania Incitti, Linda Starnes, Amy Pointon, Ryan Hicks, Benjamin E. Powell, James M.A. Turner

## Abstract

Sex chromosomes shape male (XY) - female (XX) differences in development and disease. These differences can be modelled *in vitro* by comparing XY and XX human induced pluripotent stem cells (hiPSCs). However, in this system, inter-individual autosomal variation and unstable X-dosage compensation can confound identification of sex chromosomal effects. Here, we utilise sex chromosome loss in XXY fibroblasts to generate XX and XY hiPSCs that are autosomally isogenic and exhibit stable X-dosage compensation. We also create X-monosomic (XO) hiPSCs, to investigate X-Y dosage effects. Using these autosomally isogenic lines, we examine sex differences in pluripotent stem cell expression. Transcriptional differences between XX and XY hiPSCs are surprisingly modest. However, X-haploinsufficiency induces transcriptional deregulation predominantly affecting autosomes. This effect is mediated by Y-genes with broad housekeeping functions that have X-homologues escaping X-inactivation. Our isogenic hiPSC lines provide a resource for exploring sex chromosome effects on development and disease *in vitro*.

## Introduction

Sexual dimorphisms are abundant in mammals. Historically, these were attributed to the distinct sex hormone milieu of males (XY) and females (XX). However, recent literature shows that sex chromosome complement can directly impact phenotypes in both healthy and disease state (Blencowe et al., 2022; Arnold et al., 2024; Aguet et al., 2020; Naqvi et al., 2019; Snell & Turner, 2018). For instance, two X chromosomes predispose women to autoimmune disease (Brown et al., 2022), while Y-chromosome loss in men is associated with cardiovascular disease, neurodegeneration, and cancer (Gutiérrez-Hurtado et al., 2024). Identifying how sex chromosome genes directly impact development and disease susceptibility in the absence of sex hormones represents a major challenge in biology and personal medicine.

Sex differences can be assayed *in vitro* by comparing XY and XX human induced pluripotent stem cells (hiPSCs) and their differentiated derivatives. However, two challenges are often encountered. First, since existing XY and XX hiPSC lines are derived from different individuals, they also carry differences in autosomal DNA sequence, which confound dissection of sex chromosome-derived effects. Second, XX hiPSCs often exhibit instability in X-dosage compensation, leading to gene expression artefacts (Mekhoubad et al., 2012; Tomoda et al., 2012; Bansal et al., 2021; Pasque and Plath, 2015; Bar et al., 2019; Raposo et al., 2024; Brenes et al., 2021; Topa et al., 2024). In mammals, X-dosage compensation is achieved by the collaboration of two processes. X-upregulation balances expression between the X and the autosomes, while X-inactivation, mediated by the non-coding RNA *XIST*, balances X-expression between the sexes (Disteche, 2016). Together, these processes result in an X-to-autosome (X:A) ratio close to 1.0. Previous work has shown that XX hiPSCs are prone to X-chromosome erosion, in which *XIST* expression is lost from the inactive X, and previously silent X-genes are consequently reactivated (Mekhoubad et al., 2012; Tomoda et al., 2012; Bansal et al., 2021; Pasque and Plath, 2015; Bar et al., 2019; Raposo et al., 2024; Brenes et al., 2021; Topa et al., 2024). An ideal hiPSC resource for dissecting sex differences must therefore comprise XX and XY hiPSCs that are both autosomally isogenic and exhibit stable X-dosage compensation.

Autosomally isogenic hiPSCs have been generated previously, providing important insights into the impact of sex chromosome complement on pluripotent stem cell transcription. One study reported generation of autosomally isogenic XY and XXY hiPSCs from non-mosaic XXY (Klinefelter syndrome) fibroblasts, but XX hiPSCs were not derived (Fiacco et al., 2021). Another study generated XXY, XY and XX autosomally isogenic hiPSCs, but the status of X-dosage compensation was not assayed in detail (Waldhorn et al., 2022). Further reports have compared either XX or XY cells to autosomally isogenic XO (Turner syndrome) cells; but in these instances, the parental XX cells were not isogenic to the XY cells (Ahern et al., 2021; Urbach and Benvenisty, 2009).

In this study, we report the generation of XX and XY hiPSCs that are autosomally isogenic and exhibit stable X-dosage compensation. Using our cell lines, we show that XX and XY hiPSCs exhibit surprisingly similar gene expression profiles. By generating XO hiPSCs, we also show that the Y chromosome and inactive X chromosome exert considerable effects on pluripotent stem cell gene expression.

## Results

### Generation and characterisation of autosomally isogenic XXY, XX, XY and XO hiPSCs

To generate hiPSCs clones with distinct sex chromosome complements but identical autosomes, we subjected non-mosaic XXY fibroblasts to reprogramming, which promotes sex chromosome loss (Hirota et al., 2017). We used integration-free reprogramming to avoid potential transgene expression artefacts (Beltran et al., 2020). Y-chromosome loss gave rise to XX hiPSCs, while X-chromosome loss generated XY hiPSCs (Figure 1A). Subsequently, we used CRISPR-Cas9 mediated Y-chromosome elimination to generate XO from XY hiPSCs (Figure 1A; Adikusuma et al., 2017). We achieved this using a guide RNA that targets a repeat region on the Y-chromosome centromere (Figure 1B, Figure S1A). In the resulting XO cells, off-target effects were not observed (Figure S1B).

**Figure 1.**
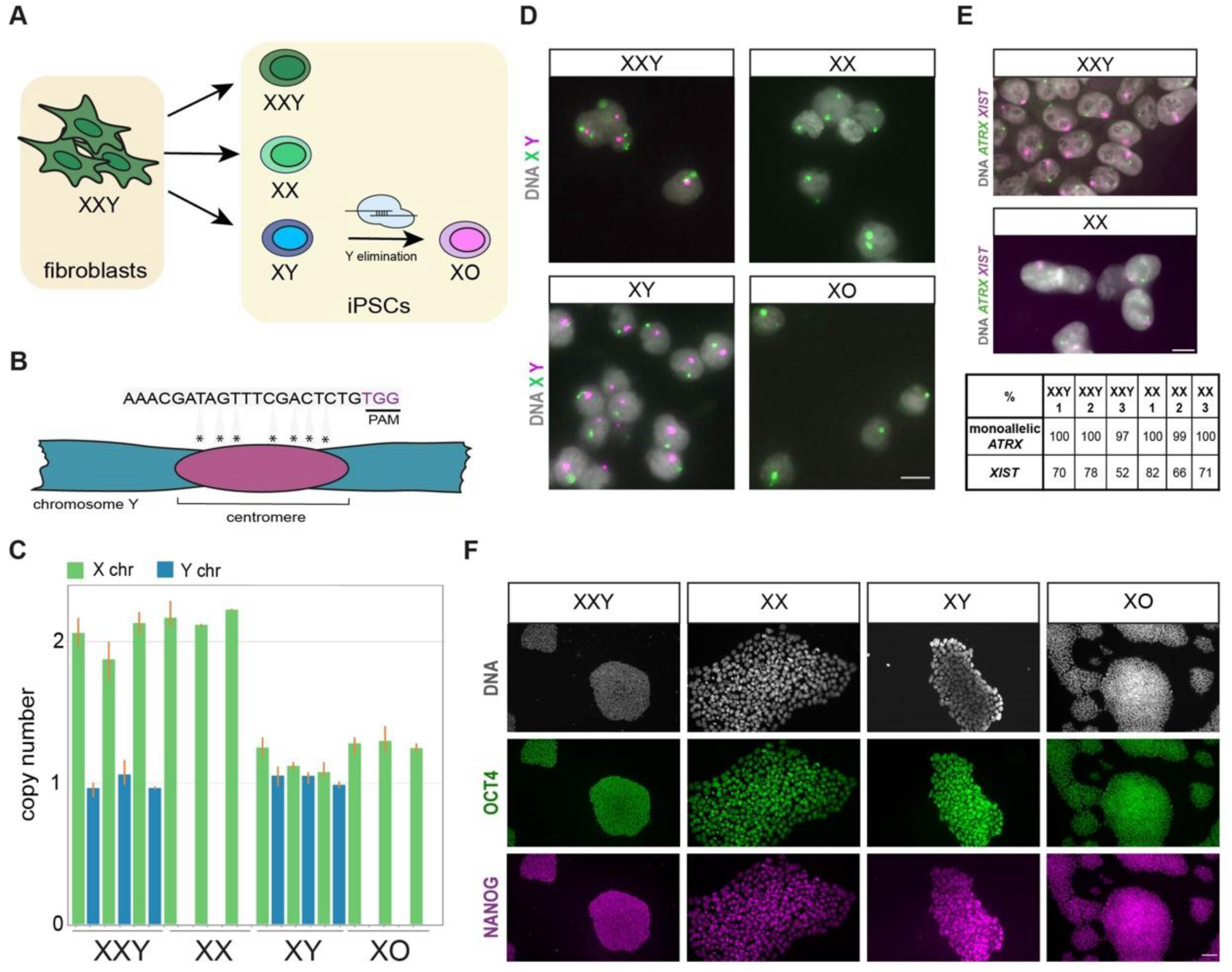
Autosomally isogenic hiPSC generation and characterisation. A) Schematic view of hiPSCs generation through XXY fibroblast reprogramming and Y-elimination via CRISPR-Cas9. B) Schematic view of seven sgRNA guide target sites on the Y centromere. C) QPCR copy number variation analysis using TaqMan *AR* (X chromosome) and *SRY* (Y chromosome) gene probes in hiPSC clones. D) Karyotype of isogenic XXY clones (n=129), XX (n=74), XY (n=119), XO (n=88) hiPSCs by X and Y DNA-FISH. The Y chromosome was present in 90%, 97%, 88%, 92%, 94% and 89% of XXY 1 (n=129), XXY 2 (n=103), XXY 3 (n=67), XY 1 (n=119), XY 2 (n=250) and XY 3 (n=119) lines, respectively. E) Representative *XIST* and *ATRX* RNA FISH analysis shows *XIST-*positive cells present in 78% and 82% of XXY 2 and XX 1 respectively. Table below shows percentage of *XIST* positive cells in XXY 1 (n=64), XXY 2 (n=71), XXY 3 (n=60), XX 1 (n=57), XX 2 (n=298), XX 3 (n=84) hiPSCs. F) OCT4 and NANOG protein expression in hiPSCs. The brightness has been increased in XXY and XO images to improve visualisation. Scale bar in D and E = 10µm, in F = 100µm.

Copy number variation qPCR and DNA FISH were used to establish the sex chromosome complement of our XXY, XX, XY and XO hiPSCs (Figure 1C, D). In addition, whole-genome sequencing was employed to confirm that they were autosomally euploid (Figure S1C). SNP analysis verified that all XY clones derived from XXY fibroblasts carried the same parental X chromosome (data not shown to maintain donor confidentiality). We screened all XXY and XX hiPSC clones for X-chromosome erosion, using a previously described approach combining RNA-FISH for *XIST* and the X-inactivated gene *ATRX* (Tchieu et al., 2010), only retaining lines which exhibited *XIST* clouds and monoallelic *ATRX* expression (Figure 1E). Additionally, we confirmed that the Y chromosome was retained in our XXY and XY hiPSC lines (see Figure 1D and legend). We ultimately generated twelve autosomally isogenic hiPSC lines, comprising triplicates for each genotype. All hiPSC lines exhibited expression of pluripotency markers POU5F1 and NANOG, as assessed by immunofluorescence (Figure 1F), as well as competence to differentiate to endoderm, mesoderm and ectoderm lineages (Figure S1D).

### Minimal gene expression differences between XX and XY iPSCs

To assay sex chromosome effects, we performed bulk RNA-seq on our hiPSC lines. As expected, Principal Component Analysis (PCA) focusing on sex-linked genes showed that the lines segregated according to their sex chromosome complement (XXY, XX, XY and XO; Figure 2A). We then repeated the PCA including all remaining genes. We found that XXY, XX and XY hiPSCs were more closely clustered to each other than to the XO hiPSCs (Figure 2B). These findings demonstrate that sex chromosomes impact autosomal genes in *trans*, and that X-chromosome monosomy has a strong impact on autosomal gene expression (see later).

**Figure 2.**
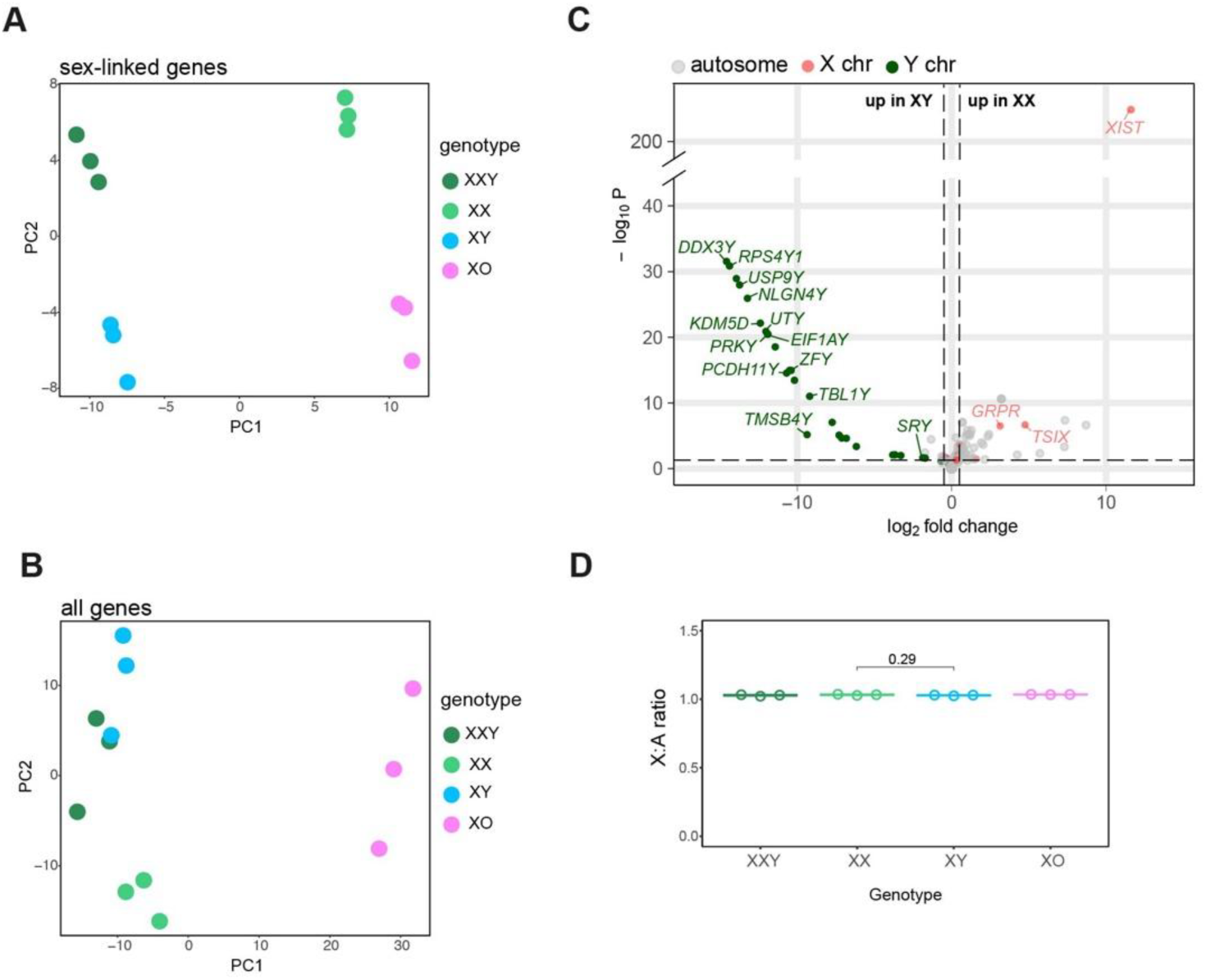
Overview of sex chromosome effects on the hiPSC transcriptional landscape. A) PCA of hiPSCs based on sex-linked gene expression. B) PCA of hiPSCs based on expression of all genes. C) Volcano plot of gene expression between XX and XY hiPSCs (note split Y axis). Horizontal dashed line represents a transformed adjusted p value of 0.05. Vertical dashed lines represent log_2_FC = 0.5. D) X:A ratio in hiPSCs.

We next examined differentially expressed (DE) genes between XX and XY hiPSCs. We observed a modest effect, with only 82 genes differentially expressed (log_2_FC0.5, padj<0.05; Figure 2C), indicating that the transcriptomes of XX and XY hiPSCs are similar (Table S1A). This finding demonstrates that the Y chromosome and inactive X chromosome have a similar impact on the transcriptome of XY and XX hiPSCs, respectively. Among the 82 DE genes, 33 were sex-linked. XX hiPSCs exhibited higher expression of *XIST, TSIX*, and *GRPR*. In contrast, most upregulated genes XY hiPSCs (87%) were Y-linked. The number of Y-genes expressed in XY hiPSCs comprised 48% of the total protein-coding gene families on the Y chromosome (13 out of 27 genes; Figure 2C). These genes included the testis-determining factor *SRY*, ware enriched for regulatory functions, e.g., chromatin modification, RNA stability and translation, and have been shown to exhibit strong purifying selection during evolution (Bellott et al., 2014; Cortez et al., 2014).

The finding that our XX and XY hiPSCs exhibited few DE genes indicated that in both genotypes X-dosage compensation was complete. We confirmed this using analysis of X:A ratios. In both the XX and XY hiPSCs, the X:A ratio was close to 1.0, demonstrating that they were indeed fully X-dosage compensated (Figure 2D). This finding also supported our earlier *ATRX* RNA-FISH analysis which showed that X-inactivation is robust in our XX hiPSCs (Figure 1E). Given the modest differences in gene expression, gene ontology analysis revealed only a few pathways differentially impacted by sex chromosome complement, with most reflecting the known functions of Y-genes, e.g., in chromatin remodelling (Figure S2A; Table S1B).

A previous study observed 313 DE genes between autosomally isogenic XX and XY hiPSCs, when employing a padj<0.1 (Waldhorn et al., 2022). When we applied the same criteria to our dataset, we observed only 162 DE genes. The overlap in DE genes between these two datasets was minimal, principally comprising Y-genes and *XIST* (Figure S3C). To understand the differences in autosomal gene expression, we compared the two datasets by PCA and Pearson correlation. PCA revealed that our XX and XY replicates (Figure 2A) showed lower gene expression variation and higher Pearson’s correlation coefficients than the Waldhorn replicates (Figure S3A and S3B).

### Sex chromosome monosomy significantly impacts hiPSC gene expression

We addressed the extent to which sex chromosomes aneuploidy impacts gene expression. Both XXY and XO hiPSCs exhibited X:A ratios that were comparable to XX and XY hiPSCs (Figure 2D). Our PCA revealed that the gene expression profile of XXY hiPSCs was similar to XX and XY hiPSCs, while the gene expression profile of XO hiPSCs was more distinct (Figure 2B). This conclusion was supported by DE analysis. We identified 63 DE genes between XXY and XX hiPSCs, 41% of which were Y-linked, and 122 DE genes between XXY and XY hiPSCs, 7% (8/122) of which were known X-inactivation escapees (Figure 3A; Table S1C-D). In comparison, a significantly larger number of genes were differentially expressed in the XO versus XY or XX hiPSC comparisons (Figure 3A; Table S1E). We found 2483 DE genes between the XY and XO hiPSCs, indicating a strong influence of the Y chromosome on the hiPSC transcriptome. In addition, 2365 genes were differentially expressed between the XX and XO hiPSCs (Table S1F), indicating a marked impact of the inactive X chromosome on the hiPSC transcriptome. Importantly, in both cases, the majority (95%) of DE genes were autosomal (Figure 3A). We conclude that X-chromosome monosomy causes significant transcriptional deregulation, and that sex chromosomes act *in trans* to modify autosomal gene expression.

**Figure 3.**
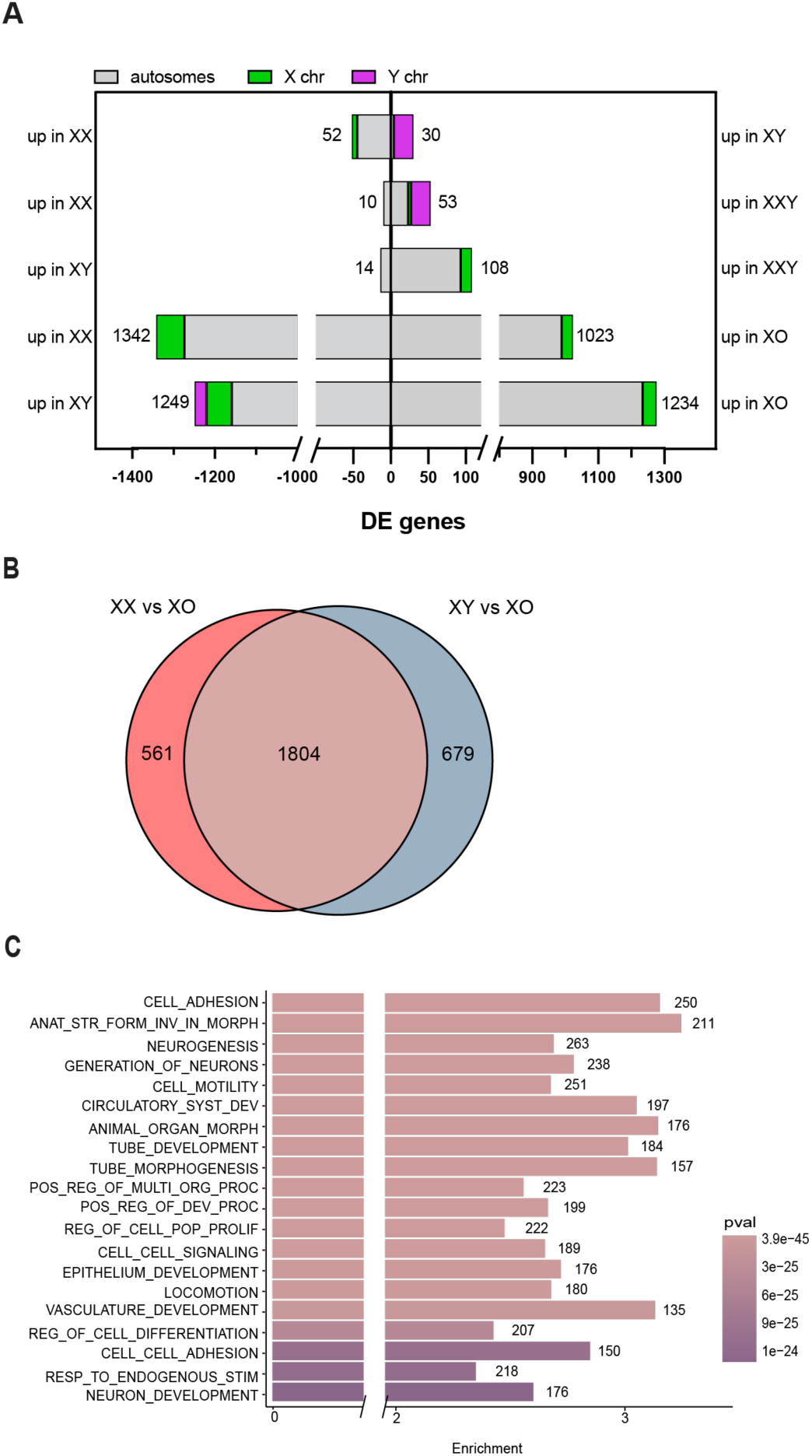
Effect of sex chromosome aneuploidy on the hiPSC transcriptional landscape. A) Bar plot of DE (FC>0.5) genes between all generated isogenic hiPSCs lines. B) Overlap of DE genes between XX versus XO and XY versus XO comparisons. C) Functional enrichment analysis of overlapping DE genes from B).

### The Y chromosome and inactive X chromosome have shared targets in hiPSCs

Our findings revealed that absence of the Y chromosome or the inactive X chromosome significantly impacted gene expression in hiPSCs. Previous work found that the Y chromosome and inactive X chromosome similarly influence transcription in human somatic cells, and that this effect is mediated by Y-genes with X-homologues that escape X-inactivation (San Roman et al., 2023; San Roman et al., 2024). Our earlier XX versus XY hiPSC comparison indicated that a similar phenomenon is conserved in hiPSCs. Interestingly ten of the thirteen Y genes we found expressed in hiPSCs (Figure 2C) have X-encoded homologues that are known to escape X-inactivation (*DDX3X, KDM5C, RPS4X, USP9X, NLGN4X, UTX, PRKX, EIF1AX, TBL1X* and *ZFX;* Bellott et al., 2014; Berletch et al., 2011). To further resolve whether the Y and inactive X have overlapping functions in hiPSCs, we looked for commonality in DE genes between the XY versus XO and XX versus XO comparisons. The degree of overlap was highly significant, comprising 73% of all XY-XO and 76% of all XX-XO DE genes (Figure 3B; Table S1G). Gene ontology analysis identified multiple cellular pathways impacted by both Y and inactive X chromosomes, including motility, adhesion and neurogenesis (Figure 3C; Table 1SH). Additionally, we detected pathways that were uniquely altered in either the XY versus XO comparison or the XX versus XO comparison (Figure S4A, B; Table S1I-L). These findings suggest that X-Y homologues act as dosage-sensitive regulators of the hiPSC transcriptome.

## Discussion

Sex chromosomes play a crucial role in male-female differences in health and disease, yet there is a lack of *in vitro* systems to fully investigate these effects (Lay Kodama and Li Gan, 2019; Miquel et al., 2023; Wilson, 2021; Wiese et al., 2023). To address this gap, we present a set of XX and XY hiPSCs that are autosomally isogenic and exhibit stable X-dosage compensation. Using these lines we find, in contrast to a previous study (Waldhorn et al., 2022), that male-female differences in hiPSC transcription are modest. Differences in reprograming methodology or source cells may partly explain this finding. However, the lower replicate variability in both our XX and XY hiPSC lines suggest that they represent an improved model for future studies of sex differences. By differentiating our autosomally isogenic hiPSCs to specific cell type in 2D or 3D manner, one could investigate sex chromosome effects on health and disease phenotypes, and test potential therapeutic targets associated with sex-linked genes (Qi et al., 2023; Rabeling & Goolam, 2023).

Dissecting the impact of sex chromosomes on sex differences can be challenging. Some effects are attributable to the presence or absence of the Y chromosome, others to X-dosage (one in males, two in females), or X-imprinting (a maternal X in males, versus a maternal and a paternal X in females; Snell and Turner, 2018). Our XO hiPSCs represent a useful tool to deconvolve Y- and X-dosage effects. By comparing transcriptomes, we show that both the Y chromosome and the inactive X chromosome regulate the hiPSC transcriptome, a phenomenon that has also been described in other cell types (San Roman et al., 2024). This function is likely mediated by deeply conserved, dosage-sensitive X-Y gene pairs. Our work adds to growing evidence that the phenotypes associated with Turner syndrome result from haploinsufficiency for these ancient X-Y homologues (Trolle et al., 2016; Bellott et al., 2014; Cortez et al., 2014).

Finally, we show that X-monosomy causes significant transcriptional deregulation. This observation may be of relevance to understanding why Turner syndrome is associated with a high rate of miscarriage (Monney et al, 2000; Gravholt et al., 1996). Our XO hiPSCs will complement others (Ahern et al., 2021; Urbach and Benvenisty, 2009) as a resource with which to identify the molecular basis of Turner syndrome phenotypes in differentiated cells and tissues.

---

**Supplementary Figure 1.**
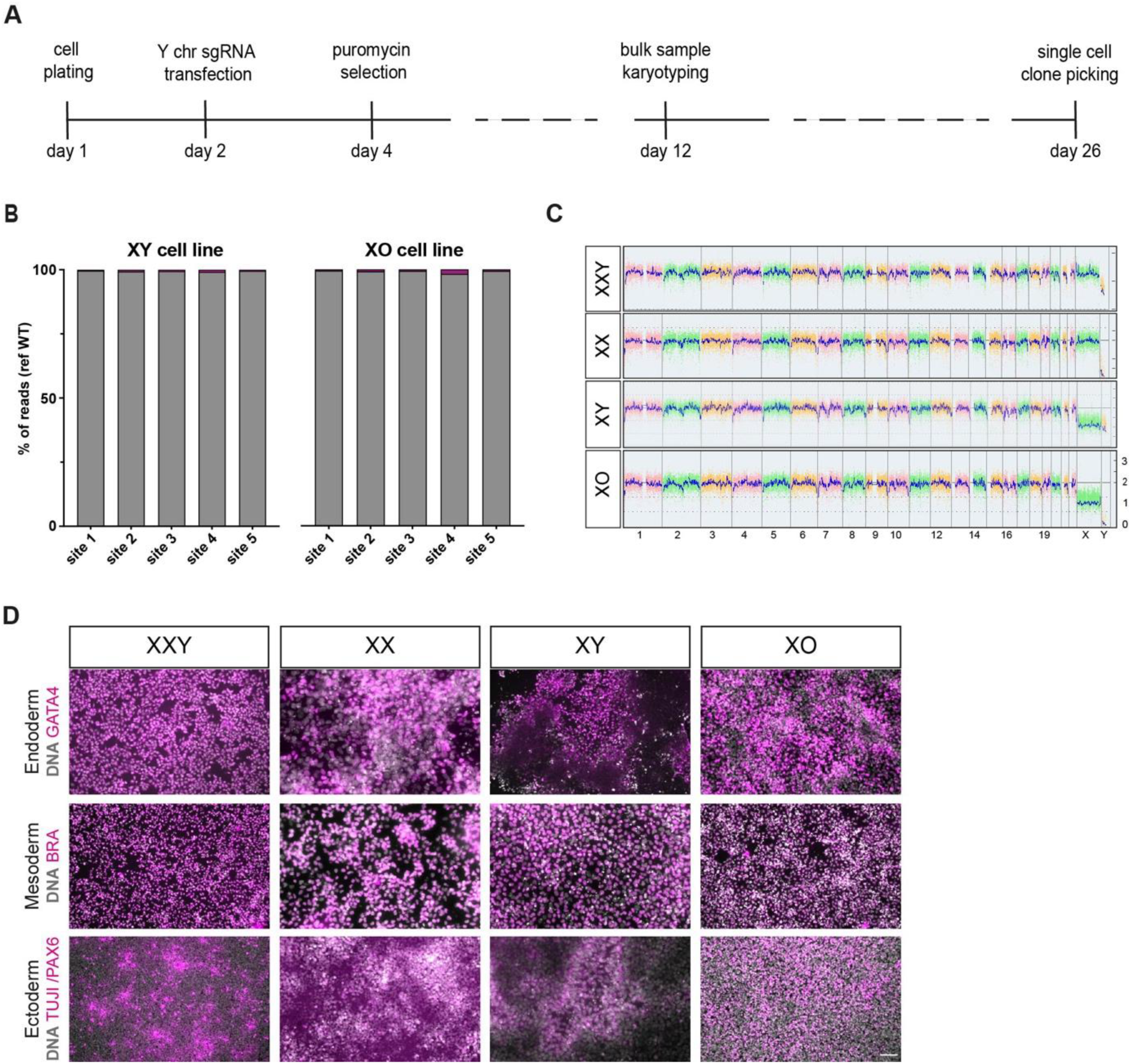
Characterisation of hiPSC lines. A) Schematic view of Y-chromosome elimination to generate XO hiPSCs. B) Analysis of Miseq reads on XY and XO hiPSCs focusing on top five off-target sites located in chr4, chr7, chr8, chr11 and chr22. C) Karyostat analysis of hiPSCs. D) Direct differentiation of hiPSCs. Scale bar in all images = 100µm.

**Supplementary Figure 2.**
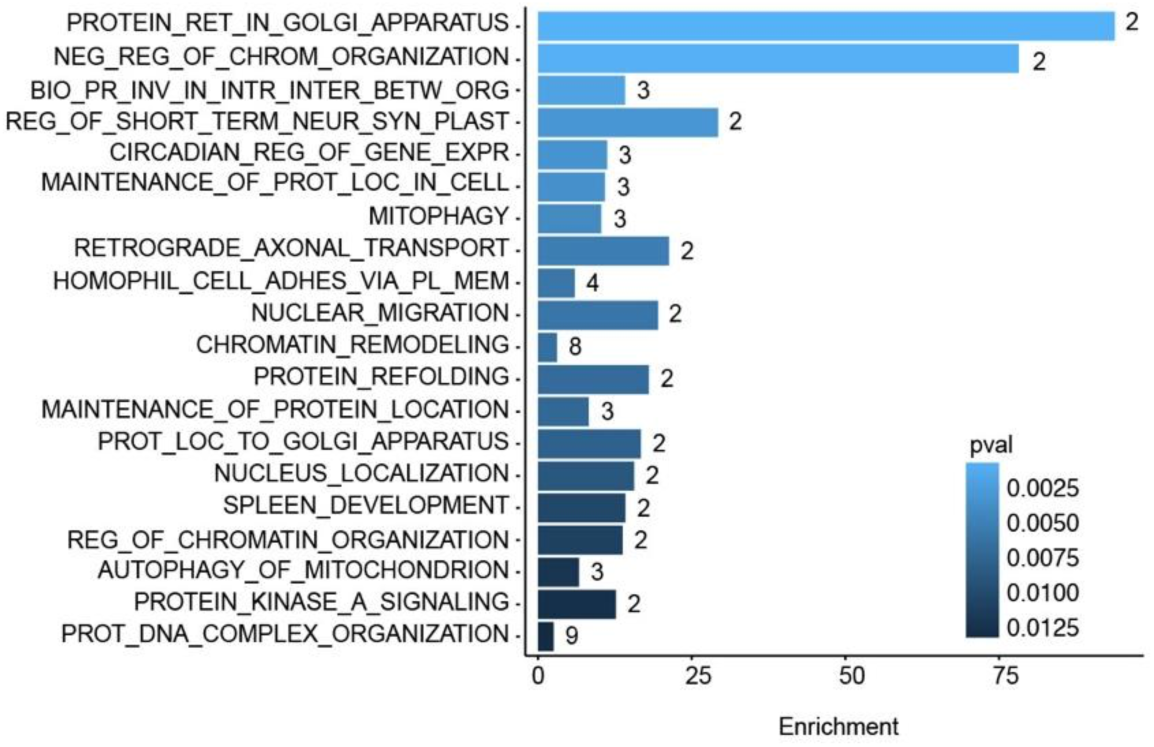
Functional enrichment analysis between XX and XY hiPSCs. Functional enrichment analysis of XX versus XY DE genes.

**Supplementary Figure 3.**
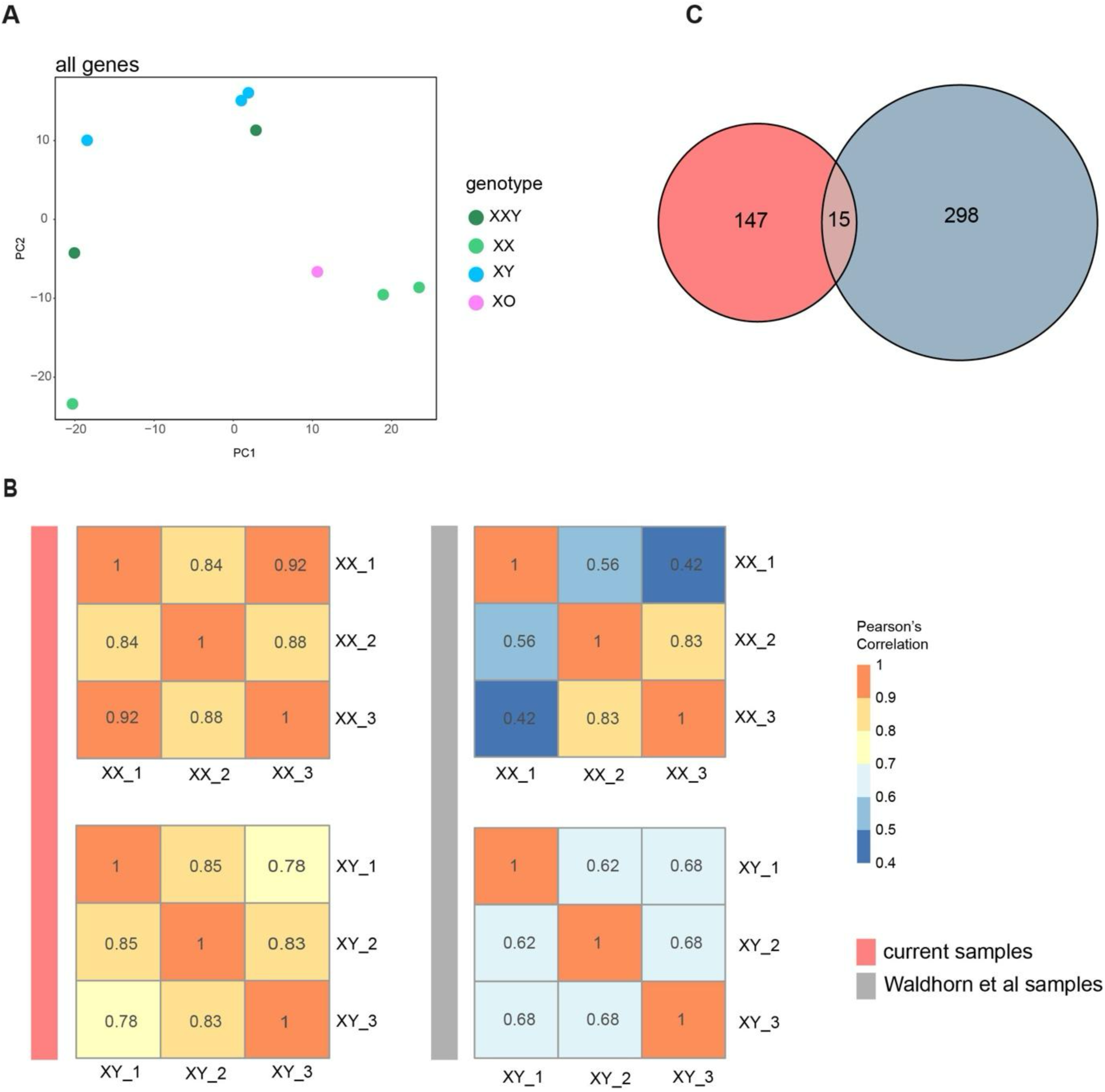
Comparison of generated hiPSCs transcriptional landscape between Waldhorn et al. Generated isogenic hiPSCs. A) PCA plot of the top 500 most variable genes for the autosomally isogenic iPSCs set generated in Waldhorn study. B) Pearson’s correlation of the top 500 most variable genes in XX and XY autosomally isogenic iPSCs samples. Sample correlations from this study are on the left, Waldhorn study – on the right. C) Euler diagram of XX vs XY DE genes between this (pink circle) and Waldhorn (grey circle) study.

**Supplementary Figure 4.**
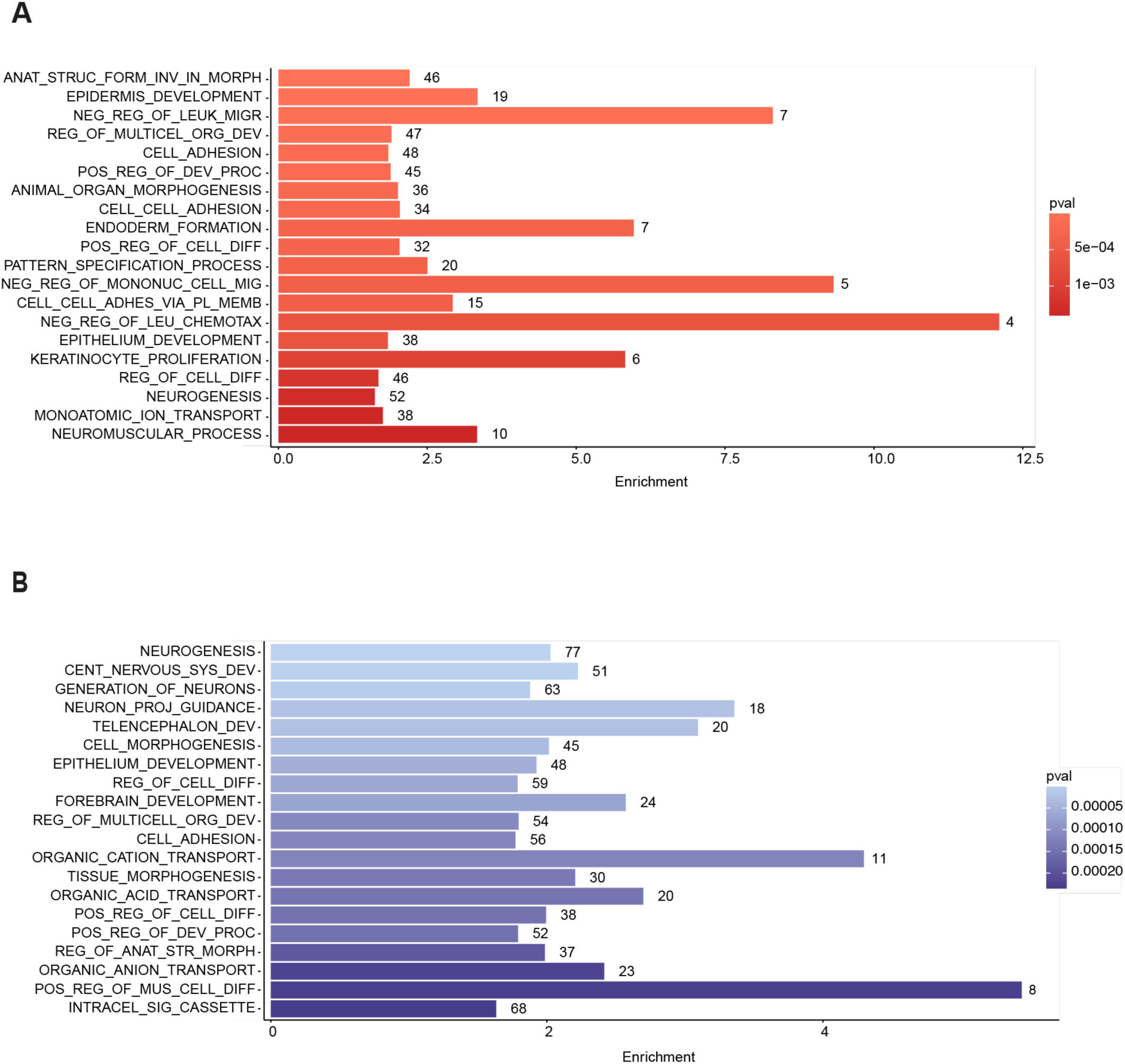
Functional enrichment analysis of DE genes between XO, XX and XY hiPSCs. A) Enrichment analysis of unique DE genes in the XX versus XO comparison. B) Enrichment analysis of unique DE genes between in the XY versus XO comparison.

## Methods

### Human fibroblast culture

GM03102 (XXY) fibroblasts were obtained from Coriell Institute for Medical Research. Fibroblasts were cultured in fibroblast medium containing Advanced DMEM/F12 medium (Thermofisher Scientific, 12634010) supplemented with 20% fetal bovine serum (Biosera, cat.no FB-1001/500).

### Human iPSCs generation and maintenance

Dermal fibroblasts were thawed at passage 6 and seeded at a density of 5×10^4^/well in 3 wells of a 6-well plate coated with 1% gelatin. Cells were plated in fibroblast medium and cultured for 48 h in 37 °C, 5% CO_2_ and 21% O_2._ The CytoTune2.0 (Thermofisher scientific, A16517) was used to reprogram the cells following kit’s recommendation. Briefly, on the first day of reprogramming, day 0, the medium was changed and complemented with sendai virus particles using 5:5:3 ratio. At day 7 cells were replated on vitronectin XF (StemCell Technologies, 7180) coated wells and at day 8 medium was changed to Essential 8 (Thermofisher Scientific, A1517001) or StemFlex medium (Thermofisher Scientific, A3349401). iPSC-like colonies started to appear 10-18 days following the transfection; they were manually picked and transferred to feeder-free conditions in vitronectin XF-coated 6-well plates with E8 or Stem Flex medium containing 10 µM Y-27362 (Biotechne, 1254) respectively. The medium was changed after 24 h. Colonies were expanded by washing with DPBS (Gibco, 14190-144) and splitting at a 1:3 to 1:6 using versene (Thermofisher Scientific, 15040066) and single cell cloning was used to select cells with sex chromosome loss. We have isolated five male clones and 8 female clones from eight hundred and thirteen single cell clones. In this study we use XXY_1, XXY_2, XXY_3, XX_1, XX_2, XX_3, XY_1, XY_2, XY_3, XO_1, XO_2 and XO_3 cell lines. Cell lines were dissociated with versene and collected in KOSR media (Thermofisher Scientific, 10828028) containing 10% DMSO (Merck, D2650) with cell lifters (Fisherbrand, 08-100-241). The working bank cells were thawed using 10 µM Y-27362.

### Short tandem repeat (STR) profiling

The Cell Services, a Science Technology Platform (STP) within the Francis Crick Institute, performed the STR profiling on DNAs from the parental fibroblasts sample and iPSC lines using the Powerplex 16 HS System (Promega, DC2101). Analysis done using Applied Biosystems 3500XL genetic analyser. All lines were sent regularly for STR profiling (every eight passages) since reprogramming started.

### Mycoplasma detection test

The Cell Services Platforn (STP) at the Francis Crick Institute confirmed the absence of mycoplasma contamination using the Phoenix DX mycoplasma mix (Procomcure, PCCSKU15209) for PCR amplification. Cells were regularly sent for Mycoplasma testing (every eight passages) since reprogramming started.

### Y chromosome sgRNA generation

Single guide RNA AAACGATAGTTTCGACTCTGTGG was designed using publicly available CRISPOR tool (Haeussler et al., 2016). The centromere region 10350000-10450000, unique to Y chromosome, was tested and a single sgRNA with predicted high target and low off target activity was selected. Following a published protocol, oligonucleotides with BbsI tails were annealed and ligated into the relevant vector (Ran et al., 2013). The sgRNA oligonucleotide was sourced from Integrated DNA Technologies.

### Primer design

All primer pairs used in this study were designed using the publicly available tool Primer3 (http://bioinfo.ut.ee/primer3/). All PCR amplifications were carried out using Q5 High-Fidelity DNA polymerase (NEB) at recommended Q5 thermocycling conditions. To amplify off target regions for MiSeq analysis, primers were designed using Primer3 and extended to contain MiSeq adaptor sequences (see relevant section). All primer sequences are listed in Table S2.

### Transfection assay

Human iPSCs were maintained in Advanced E8 or StemFlex medium and plated as single cells a day before transfection. On day 1, cells were transfected with the Cas9-eGFP vector plasmid containing sgRNA targeting Y chromosome, Optimem (Gibco, 31985-062) and FuGeneHD buffer (Promega, E2311). Ratios were used following manufacturer’s instructions. Targeted hiPSC clones were selected by adding puromycin (0.5-1 ug/ml) when they’ve reached 80% confluency on day 2 or day 3 for 24-48 hours. Targeted colonies appeared by day 6-12. Transfection efficiency is dependent on cell line and was recorded between 5 % - 45 %. In this study we use XY_1, XY_2 and XY_3 cell lines, passage numbers between P10 - P15.

### Immunofluorescence assay

We used immunostaining assay to evaluate pluripotency and germ layer precursors potential. IPSCs (passage between P5 - P26) were fixed with 4% paraformaldehyde in DPBS (Thermofisher Scientific, 14190094) for 1h at RT, then washed three times with PBS-A. Blocking and permeabilization was achieved by incubation with dPBS with 3% bovine serum albumin (Merck, A9647) and 1% Triton X- 100 (Merck, T8787) for 1h at room temperature. Primary antibodies were diluted in blocking solution and incubated overnight at 4 C. Following incubation, the cells underwent 2 x 5 min washes with PBS-A. Secondary antibodies were diluted 1:200 in 3% BSA and incubated for 2 h at room temperature. After 3 x 5 min washes in PBS-A, 5 μg/ml DAPI (Merck, D9542) was added for 5 mins during the middle wash to perform nuclear staining. The used primary antibodies are listed in Table S3.

### Direct differentiation assay

The STEMdiff™ Trilineage Differentiation Kit (StemCell Technologies, 5230) was used to differentiate cells into three germ lineage precursors. The assay was performed as per kit instructions. We evaluated the differentiation potential of all lines by immunostaining for lineage-specific markers on day 5 (Mesoderm and Endoderm) and day 7 (Ectoderm). Cell lines’ passage numbers were between P10 - P26 for this experiment.

### Genomic DNA isolation and copy number variation assay

Cells were collected from the well and pelleted down after centrifugation at x300g for 4min. Cell pellet was resuspended in 200ul of dPBS. Invitrogen PureLink Genomic DNA kit (Invitrogen, K1820-02) was used to isolate DNA following kit’s instructions. Isolated DNA concentration was measured using Nanodrop 2000c device (Thermo Scientific).

TaqMan Copy Number Assays (Thermofisher Scientific) were used with probes for SRY on the Y chromosome and AR on the X chromosomes (Hs01026408_cn and Hs04511283_ cn respectively). Each sample was subjected to duplex TaqMan qRT-PCR according to Applied Biosystems protocol. All reactions were run in quadruplicates. Data were analysed using CopyCaller version 2.1 software (Applied Biosystems). Previously established XX and XY human iPSCs were used as control samples.

### DNA-FISH

Centromeric probes for X and Y chromosome, MetaSystems XA X/Y (Pisces Scientific, D-5608-100-OG), were used to assess X and Y chromosome number in cells. Cell lines of passage between P4 - P10 were attached on glass slides by growing cells on slides (fibroblasts) or air-drying dissociated cells on slides (iPSCs). Cells were fixed in 4% paraformaldehyde in phosphate-buffered saline (PBS) at room temperature for 10 minutes and permeabilized in 0.05% Triton X-100 in PBS at room temperature for 10 minutes. Slides were washed in 2x SSC at room temperature and 75 °C and denatured in 70% formamide in 2x SSC. After dehydration in an ethanol series (70, 80, 95, 100%), air-dried specimens were hybridized with the denatured probe in hybridization buffer (2x SSC, 50% formamide, 25% dextran sulphate, 5 mg/ml bovine serum albumin, 1mM Vanadyl Ribonucleoside) at 37 °C for overnight. After wash in 2x SSC at 45 °C, 0.1x SSC at 60°C, and 4xSSC, 0.1 % Tween20 at room temperature, slides were mounted in Vectashield with DAPI (2BScientific, H-1200-10).

### RNA-FISH

Cells of passage P10 - P15 were grown on slides or dissociated and air-dried on the cooled slides. Cold permabilisation buffer (0.5%Triton X-100, 2 mM vanadyl ribonucleoside in PBS-A) was added on the slides for 10min following fixation with cold 4% PFA. After removing the fixation solution, slides were rinsed in ice-cold PBS-A, dehydrated through ice-cold 70%, 80%, 95% and 100% ethanol for 5 minutes each, and air dried. BAC XIST RP11-13M9 (CHORI) and BAC ATRX RP11-1145J4 (CHORI) were labelled using Nick Translation Direct Labelling Kit (Abbott Laboratories, 07J00-01) according to manufacturer’s instructions, and using fluorescent nucleotides (for XIST spectrum orange – dUTP; Abbott, 02N32-050, for ATRX spectrum green – dUTP; Abbott, 02N33-050). Cells were hybridized with a denatured mix of probes along with 1 µg salmon sperm DNA in hybridization buffer (50% formamide, 25% dextran sulphate, 5 mg/ml BSA, 1mM Vanadyl Ribonucleoside complex in 2X SSC) at 37°C overnight in a humid chamber. Stringency washes were performed on a hot plate, three times for 5 mins in 50% formamide, 1X SSC (pH 7.2-7.4) pre-heated to 45°C, and three times for 5 mins in 2X SSC (pH 7-7.2) pre heated to 45°C. Cells were stained with DAPI in 2xSSC (1ug/ml) for 10 mins at room temp, rinsed once with 2X SSC, and mounted in Vectashield with DAPI and stored at -20°C.

### Karyotyping assay

Chromosomal microarray was performed using genomic DNA of the iPSC lines (P10 - P15), Thermo Scientific (USA) performed the KaryoStat assay (Thermo Scientific, USA), an array comparative genomic hybridization (CGH).

### MiSeq high throughput sequencing

Five off-target sites in chr 4, chr7, chr11 and chr22 were tested by amplifying the regions using MiSeq PCR primers (Table S2.) in a total volume of 25 μl (12.5 μl NEB Q5 High-Fidelity Master Mix, 5 mM each primer). Correct PCR amplification was confirmed by gel electrophoresis. Resultant PCR amplicons were purified using beads AMPure XP (Beckman coulter, A63881) and resuspended in 15 μl nuclease-free water and prepared according to the Illumina MiSeq library prep manufacturer’s instructions (Nextera Index Kit V2). Libraries are quantified using Promega Quantifluor reagents, plate reader and Agilent Tapestation. MiSeq libraries are pooled by concentration and sequenced on the Illumina MiSeq-Nano platform with a PE 250bp run configuration on a Nano flowcell. On average samples receive 2,000-5,000 reads each.

Resultant reads were demultiplexed and fastq files were collapsed using FastX Toolkit (v0.0.13) [https://github.com/agordon/fastx_toolkit]. To assess the rate of indel-production by CRISPR-Cas9, the reads were aligned to the human reference genome hg38 with the Burrows-Wheeler Alignment tool (BWA, v0.7.170) (Li and Durbin, 2009) using the mem algorithm with default settings and then analysed using the R package CrispRVariants (v1.14.0) (Lindsay et al., 2016). The proportion of wild type reads in off target sites were calculated by dividing the number of reads matched to reference genome by sum of reference reads and all reads containing single nucleotide variants ± 20bp away from the sgRNA binding site.

### Microscopy and Image Analysis

Immunostainings for pluripotency and trilineage markers, DNA and RNA-FISH images were acquired using the Whitefield Imaging system AxioImager.M1 (Zeiss) and MicroManager (2.0) using 10X and 40X oil objectives. 150 (FITC), 400 (Cy3), and 400 (Cy5) laser power were used for pluripotency markers; 300 (FITC), 300 (Cy3), and 300 (Cy5) used for trilineage germ layer markers; and 400 (FITC), and 400 (Cy3) were used for DNA/RNA-FISH markers. At least 50 single cells spread over 3 to 8 fields were acquired for each cell line. The number of nuclei/fields was defined by DAPI-positive cells, and the percentage of positive signals for XIST and ATRX RNA-FISH, as well as X- and Y-centromeric signals for DNA-FISH, was calculated. Cells after Images were processed and analysed using ImageJ software (v2.1).

### RNA isolation

RNA was extracted using TRI Reagent (Merck, T9424) following the chloroform-isopropanol protocol (Chomczynski and Sacchi, 2006). Briefly, the cells were pelleted and dissolved in 500ul TRI reagent. After adding 100ul of chloroform, cell lysate was vigorously shaken and span in cold centrifuge for 30min. Aqueous phase was collected and precipitated using isopropanol. The pellet was dried out and dissolved in 50ul of RNase free water.

### Bulk RNA-Seq Library Preparation and Sequencing

RNA libraries were generated using the KAPA mRNA HyperPrep Kit (Roche, KK8581) according to manufacturer’s instructions. Briefly, samples of passage P5 - P20 were normalised to 100 ng -1000 ng of RNA in a final volume of 50 ul. PolyA-tailed RNA capture was performed twice with 50 ul of capture beads at 65 °C for 2 min first and 70°C for 2 min for second capture following 20 °C for 5 min both times. Beads were washed to remove other RNA species after each capture and eluted in 20 ul of Fragment, Prime and Elute Buffer. The fragmentation reaction was run for 6 min at 94 °C for a library insert size of 200-300 bp. Fragmented RNA underwent first and second strand cDNA synthesis according to manufacturer instructions. The adaptor ligation was carried out using the KAPA Unique Dual-Indexed Adapters Kit (15μM) (Roche, KK8727) stock diluted to 1.5-7 uM. 50ul of ligation buffer and 10 ul of DNA ligase were added to 60ul of cDNA and incubated at 20°C for 15 min. SPRISelect beads 0.63 -0.7x were used to remove short fragments. Library was amplified using 20ul cDNA with 25ul of Kapa HiFi HotStart PCR master mix plus 5ul of Library Amplification Primer mix. 8-13 PCR cycles were performed as recommended by the manufacturer for a total RNA input of 100-1000 ng. Amplified libraries were purified via a SPRISelect 1x bead cleanup. The quality and fragment size distributions of the purified libraries was assessed by a 4200 TapeStation Instrument (Agilent Technologies). Libraries were pooled and sequenced on the Illumina NovaSeq6000 in PE100 configuration to an average depth of 25M paired-end reads.

### RNA-Seq data profiling

Raw RNA-seq reads were processed using the RNA-seq nf-core pipeline (v3.14). Reads were aligned to the hg38 genome using star_rsem. Specifically, XXY and XY genotypes were aligned to the standard hg38 genome while the XX and XO genotypes was aligned to a modified hg38 genome which did not include the Y chromosome sequence. The two resulting raw counts tables were merged. The raw counts were processed in R using the DESeq2 package (v1.40.2). Very lowly expressed genes were filtered out by applying a rowSums filter of >=10 to the raw counts table. Counts were normalised using the DESeqDataSetFromMatrix() and DESeq() functions, specifying “∼genotype” in the design formula. For comparisons against the Waldhorn dataset “∼batch + genotype” was specified as the design formula. Principal component analysis (PCA) plots were generated using the top 500 most variable genes, applying the vst() function for PCA visualisation. Differentially expressed genes were identified using the lfcShrink() function, specifying “ashr” as the shrinkage estimator. Differentially expressed genes (DEGs) were defined as genes which had an adjusted p-value ≤ 0.05 and log2FC ≥ 0.5. Functional enrichment of differentially expressed genes was done as described in Roman *et al*., 2024. Correlation heatmaps between XX and XY replicates was done by identifying the top 500 most variable genes in the XX or XY cohort then calculating the Pearson correlation coefficient between replicates.

### Whole Exome Sequencing

DNA libraries were prepared using 200 ng of genomic DNA, fragmented to a target size of 150 to 200 bp on the Covaris E220, as input into an Agilent SureSelect XT library preparation kit, and whole-exome capture was performed using a Human All Exon V5 capture library according to the manufacturer’s guidelines. Libraries were then multiplexed and sequenced using 100 bp paired-end reads on Illumina HiSeq 4000 to a depth of at least 50M paired-end reads per sample.

### SNP identification on X chromosome

DNA sequence data were trimmed using trim_galore, with the following parameters --paired --fastqc --gzip --retain_unpaired --three_prime_clip_R1 2 --three_prime_clip_R2 2. Reads were mapped using bwa memagainst the human reference hg38 genome using default parameters. Alignment files were converted from sam to bam format, sorted and indexed using samtools view -b, sort and index respectively. PCR doublets were marked using Picard. Variant calling was performed using gatk specifying HaplotypeCaller parameter. IGV (v2.9.4) was used to check SNPs in all transcribed X-linked genes in generated iPSC lines.

## Acknowledgments

This work was supported by AstraZeneca – Crick Collaboration (CC2052) and by Francis Crick Institute which received its core funding from Cancer Reaserach UK (CC2052), the UK Medical Research Council (CC2052) and the Wellcome Trust (CC2052). The authors thank the Francis Crick Institute Human Biology, Advanced Light Microscopy and Advanced Sequencing facilities for their contributions and expertise. We thank the members of the J.M.A.T. lab for comments and discussion on the manuscript.

## Author contributions

Study Conception and Design, J.M.A.T. and B.E.P; Investigation and Data Analysis, R.M.; Bioinformatic Data Analysis, W.V. and J.Z.; Manuscript Writing and Revision, J.M.A.T., R.M., W.V., B.E.P., L.S., T.I., J.E., R.H., and A.P; Supervision, J.M.A.T., R.H., and A.P.

## References

1. Adikusuma F, Williams N, Grutzner F, Hughes J, & Thomas P. (2017). Targeted Deletion o an Entire Chromosome Using CRIPSR/Cas9. Mol Ther., 25(*8*), 1736–1738. 10.1016/j.ymthe.2017.05.021.

2. Aguet, F., Barbeira, A. N., Bonazzola, R., Brown, A., Castel, S. E., Jo, B., Kasela, S., Kim-Hellmuth, S., Liang, Y., Oliva, M., Flynn, E. D., Parsana, P., Fresard, L., Gamazon, E. R., Hamel, A. R., He, Y., Hormozdiari, F., Mohammadi, P., Muñoz-Aguirre, M., … Volpi, S. (2020). The impact of sex on gene expression across human tissues. Science, 369(6509). 10.1126/SCIENCE.ABA3066.

3. Ahern, D. T., Bansal, P., Faustino, I., Kondaveeti, Y., Glatt-Deeley, H. R., Banda, E. C., & Pinter, S. F. (2021). Monosomy X in isogenic human iPSC-derived trophoblast model impacts expression modules preserved in human placenta. PNAS. 10.1101/2021.12.13.472325.

4. Arnold, A. P., Klein, S. L., Mccarthy, M. M., & Mogil, J. S. (2024). Male-female comparisons are powerful in biomedical research. Nature, 629, 37–40. 10.1038/d41586-024-01205-2.

5. Bansal, P., Ahern, D. T., Kondaveeti, Y., Qiu, C. W., & Pinter, S. F. (2021). Contiguous erosion of the inactive X in human pluripotency concludes with global DNA hypomethylation. Cell Reports, 35(10). 10.1016/j.celrep.2021.109215.

6. Bar, S., Seaton, L. R., Weissbein, U., Eldar-Geva, T., & Benvenisty, N. (2019). Global Characterization of X Chromosome Inactivation in Human Pluripotent Stem Cells. Cell Reports, 27(1), 20–29.e3. 10.1016/j.celrep.2019.03.019.

7. Bellott, D. W., Hughes, J. F., Skaletsky, H., Brown, L. G., Pyntikova, T., Cho, T. J., Koutseva, N., Zaghlul, S., Graves, T., Rock, S., Kremitzki, C., Fulton, R. S., Dugan, S., Ding, Y., Morton, D., Khan, Z., Lewis, L., Buhay, C., Wang, Q., … Page, D. C. (2014). Mammalian y chromosomes retain widely expressed dosage-sensitive regulators. Nature, 508(7497), 494–499. 10.1038/nature13206.

8. Beltran, A. A., Molina, S. G., & Beltran, A. S. (2020). Derivation of Induced Pluripotent Stem Cells from Human Fibroblasts Using a Non-integrative System in Feeder-free Conditions. Bio-Protocol, 10(20). 10.21769/BioProtoc.3788.

9. Berletch, J. B., Yang, F., Xu, J., Carrel, L., & Disteche, C. M. (2011). Genes that escape from X inactivation. Human Genetics, 130(2), 237–245. 10.1007/s00439-011-1011-z.

10. Blencowe, M., Chen, X., Zhao, Y., Itoh, Y., McQuillen, C. N., Han, Y., Shou, B. L., McClusky, R., Reue, K., Arnold, A. P., & Yang, X. (2022). Relative contributions of sex hormones, sex chromosomes, and gonads to sex differences in tissue gene regulation. Genome Research, 32(5), 807–824. 10.1101/gr.275965.121.

11. Brenes, A. J., Yoshikawa, H., Bensaddek, D., Mirauta, B., Seaton, D., Hukelmann, J. L., Jiang, H., Stegle, O., & Lamond, A. I. (2021). Erosion of human X chromosome inactivation causes major remodeling of the iPSC proteome. Cell Reports, 35(4). 10.1016/j.celrep.2021.109032.

12. Brown, G. J., Cañete, P. F., Wang, H., Medhavy, A., Bones, J., Roco, J. A., He, Y., Qin, Y., Cappello, J., Ellyard, J. I., Bassett, K., Shen, Q., Burgio, G., Zhang, Y., Turnbull, C., Meng, X., Wu, P., Cho, E., Miosge, L. A., … Vinuesa, C. G. (2022). TLR7 gain-of-function genetic variation causes human lupus. Nature, 605(7909), 349–356. 10.1038/s41586-022-04642-z.

13. Chomczynski, P., & Sacchi, N. (2006). The single-step method of RNA isolation by acid guanidinium thiocyanate-phenol-chloroform extraction: Twenty-something years on. Nature Protocols, 1(2), 581–585. 10.1038/nprot.2006.83.

14. Cortez, D., Marin, R., Toledo-Flores, D., Froidevaux, L., Liechti, A., Waters, P. D., Grützner, F., & Kaessmann, H. (2014). Origins and functional evolution of y chromosomes across mammals. Nature, 508(7497), 488–493. 10.1038/nature13151.

15. Disteche, C. M. (2016). Dosage compensation of the sex chromosomes and autosomes. In Seminars in Cell and Developmental Biology (Vol. 56, pp. 9–18). Academic Press. 10.1016/j.semcdb.2016.04.013.

16. Fiacco, E., Alowaysi, M., Astro, V., & Adamo, A. (2021). Generation of an iPSC cohort of isogenic iPSC lines (46-XY and 47-XXY) from a non-mosaic Klinefelter Syndrome patient (47-XXY) (KAUSTi008-A, KAUSTi008-B, KAUSTi008-C, KAUSTi008-D, KAUSTi008-E, KAUSTi008-F, KAUSTi008-G). Stem Cell Research, 50. 10.1016/j.scr.2020.102119.

17. Gravholt C.H., Juul S., Weis Naeraa R., & Hansen J. (1996). Prenatal and postnatal prevalence of Turner’s syndrome: a registry study. BMJ, 312, 16–21.

18. Gutiérrez-Hurtado, I. A., Sánchez-Méndez, A. D., Becerra-Loaiza, D. S., Rangel-Villalobos, H., Torres-Carrillo, N., Gallegos-Arreola, M. P., & Aguilar-Velázquez, J. A. (2024). Loss of the Y Chromosome: A Review of Molecular Mechanisms, Age Inference, and Implications for Men’s Health. In International journal of molecular sciences (Vol. 25, Issue 8). 10.3390/ijms25084230.

19. Haeussler, M., Schönig, K., Eckert, H., Eschstruth, A., Mianné, J., Renaud, J. B., Schneider-Maunoury, S., Shkumatava, A., Teboul, L., Kent, J., Joly, J. S., & Concordet, J. P. (2016). Evaluation of off-target and on-target scoring algorithms and integration into the guide RNA selection tool CRISPOR. Genome Biology, 17(1). 10.1186/s13059-016-1012-2.

20. Hirota, T., Ohta, H., Powell, B. E., Mahadevaiah, S. K., Ojarikre, O. A., Saitou, M., & Turner, J. M. A. (2017). Fertile offspring from sterile sex chromosome trisomic mice. Science, 357(6354), 932–935. 10.1126/science.aam9046.

21. Lau, Y. F. C. (2020). Y chromosome in health and diseases. In Cell and Bioscience (Vol. 10, Issue 1). BioMed Central Ltd. 10.1186/s13578-020-00452-w.

22. Lay Kodama, & Li Gan. (2019). Do Microglial Sex Differences Contribute to sex Differences in Neurodegenerative Diseases? Trends Mol Med. doi:10.1016/j.molmed.2019.05.001.

23. Lentini, A., Cheng, H., Noble, J. C., Papanicolaou, N., Coucoravas, C., Andrews, N., Deng, Q., Enge, M., & Reinius, B. (2022). Elastic dosage compensation by X-chromosome upregulation. Nature Communications, 13(1). 10.1038/s41467-022-29414-1.

24. Li, H., & Durbin, R. (2009). Fast and accurate short read alignment with Burrows-Wheeler transform. Bioinformatics, 25(14), 1754–1760. 10.1093/bioinformatics/btp324.

25. Lindsay, H., Burger, A., Biyong, B., Felker, A., Hess, C., Zaugg, J., Chiavacci, E., Anders, C., Jinek, M., Mosimann, C., & Robinson, M. D. (2016). CrispRVariants charts the mutation spectrum of genome engineering experiments. In Nature Biotechnology (Vol. 34, Issue 7, pp. 701–702). Nature Publishing Group. 10.1038/nbt.3628.

26. Mekhoubad, S., Bock, C., De Boer, A. S., Kiskinis, E., Meissner, A., & Eggan, K. (2012). Erosion of dosage compensation impacts human iPSC disease modeling. Cell Stem Cell, 10(5), 595–609. 10.1016/j.stem.2012.02.014.

27. Miquel, C. H., Faz-Lopez, B., & Guéry, J. C. (2023). Influence of X chromosome in sex-biased autoimmune diseases. Journal of Autoimmunity, 137. 10.1016/j.jaut.2023.102992.

28. Monney C, Pescia G, Addor MC. Le syndrome de Turner [Turner syndrome]. Schweiz Med Wochenschr. 2000 Sep 23;130(38):1339–43. French. PMID: 11064926.

29. Naqvi, S., Godfrey, A. K., Hughes, J. F., Goodheart, M. L., Mitchell, R. N., & Page, D. C. (2019). Conservation, acquisition, and functional impact of sex-biased gene expression in mammals. Science, 365(6450). 10.1126/science.aaw7317.

30. Pasque, V., & Plath, K. (2015). X chromosome reactivation in reprogramming and in development. In Current Opinion in Cell Biology (Vol. 37, pp. 75–83). Elsevier Ltd. 10.1016/j.ceb.2015.10.006.

31. Qi, M., Pang, J., Mitsiades, I., Lane, A. A., & Rheinbay, E. (2023). Loss of chromosome Y in primary tumors. Cell, 186(14), 3125–3136.e11. 10.1016/j.cell.2023.06.006.

32. Rabeling, A., & Goolam, M. (2023). Cerebral organoids as an in vitro model to study autism spectrum disorders. In Gene Therapy (Vol. 30, Issue 9, pp. 659–669). Springer Nature. 10.1038/s41434-022-00356-z.

33. Ran, F. A., Hsu, P. D., Wright, J., Agarwala, V., Scott, D. A., & Zhang, F. (2013). Genome engineering using the CRISPR-Cas9 system. Nature Protocols, 8(11), 2281–2308. 10.1038/nprot.2013.143.

34. Raposo, A. C., Caldas, P., Arez, M., Jeremias, J., Barbosa, P., Sousa-Luís, R., Água, F., Oxley, D., Mupo, A., Eckersley-Maslin, M., Casanova, M., Grosso, A. R., & da Rocha, S. T. (2024). Erosion of X-Chromosome Inactivation in female hiPSCs is heterogeneous and persists during differentiation. 10.1101/2024.03.15.585169.

35. San Roman, A. K., Godfrey, A. K., Skaletsky, H., Bellott, D. W., Groff, A. F., Harris, H. L., Blanton, L. V., Hughes, J. F., Brown, L., Phou, S., Buscetta, A., Kruszka, P., Banks, N., Dutra, A., Pak, E., Lasutschinkow, P. C., Keen, C., Davis, S. M., Tartaglia, N. R., … Page, D. C. (2023). The human inactive X chromosome modulates expression of the active X chromosome. Cell Genomics, 3(2). 10.1016/j.xgen.2023.100259.

36. San Roman, A. K., Skaletsky, H., Godfrey, A. K., Bokil, N. V., Teitz, L., Singh, I., Blanton, L. V., Bellott, D. W., Pyntikova, T., Lange, J., Koutseva, N., Hughes, J. F., Brown, L., Phou, S., Buscetta, A., Kruszka, P., Banks, N., Dutra, A., Pak, E., … Page, D. C. (2024). The human Y and inactive X chromosomes similarly modulate autosomal gene expression. Cell Genomics, 4(1). 10.1016/j.xgen.2023.100462.

37. Snell, D. M., & Turner, J. M. A. (2018). Sex Chromosome Effects on Male–Female Differences in Mammals. In Current Biology (Vol. 28, Issue 22, pp. R1313–R1324). Cell Press. 10.1016/j.cub.2018.09.018.

38. Subrini, J., & Turner, J. (2021). Y chromosome functions in mammalian spermatogenesis. 0. 10.7554/eLife.

39. Tallaksen, H. B. L., Johannsen, E. B., Just, J., Viuff, M. H., Gravholt, C. H., & Skakkebæk, A. (2023). The multi-omic landscape of sex chromosome abnormalities: current status and future directions. In Endocrine Connections (Vol. 12, Issue 9). BioScientifica Ltd. 10.1530/EC-23-0011.

40. Tchieu, J., Kuoy, E., Chin, M. H., Trinh, H., Patterson, M., Sherman, S. P., Aimiuwu, O., Lindgren, A., Hakimian, S., Zack, J. A., Clark, A. T., Pyle, A. D., Lowry, W. E., & Plath, K. (2010). Female human iPSCs retain an inactive X chromosome. Cell Stem Cell, 7(3), 329–342. 10.1016/j.stem.2010.06.024.

41. Tomoda, K., Takahashi, K., Leung, K., Okada, A., Narita, M., Yamada, N. A., Eilertson, K. E., Tsang, P., Baba, S., White, M. P., Sami, S., Srivastava, D., Conklin, B. R., Panning, B., & Yamanaka, S. (2012). Derivation conditions impact x-inactivation status in female human induced pluripotent stem cells. Cell Stem Cell, 11(1), 91–99. 10.1016/j.stem.2012.05.019.

42. Topa, H., Benoit-Pilven, C., Tukiainen, T., & Pietiläinen, O. (2024). X-chromosome inactivation in human iPSCs provides insight into X-regulated gene expression in autosomes. Genome Biology, 25(1). 10.1186/s13059-024-03286-8.

43. Trolle, C., Nielsen, M. M., Skakkebæk, A., Lamy, P., Vang, S., Hedegaard, J., Nordentoft, I., Ørntoft, T. F., Pedersen, J. S., & Gravholt, C. H. (2016). Widespread DNA hypomethylation and differential gene expression in Turner syndrome. Scientific Reports, 6. 10.1038/srep34220.

44. Urbach, A., & Benvenisty, N. (2009). Studying early lethality of 45,XO (Turner’s syndrome) embryos using human embryonic stem cells. PLoS ONE, 4(1). 10.1371/journal.pone.0004175.

45. Waldhorn, I., Turetsky, T., Steiner, D., Gil, Y., Benyamini, H., Gropp, M., & Reubinoff, B. E. (2022). Modeling sex differences in humans using isogenic induced pluripotent stem cells. Stem Cell Reports, 17(12), 2732–2744. 10.1016/j.stemcr.2022.10.017.

46. Wiese, C. B., Avetisyan, R., & Reue, K. (2023). The impact of chromosomal sex on cardiometabolic health and disease. In Trends in Endocrinology and Metabolism (Vol. 34, Issue 10, pp. 652–665). Elsevier Inc. 10.1016/j.tem.2023.07.003.

47. Wilson, M. A. (2021). The y chromosome and its impact on health and disease. In Human Molecular Genetics (Vol. 30, Issue 2, pp. R296–R300). Oxford University Press. 10.1093/hmg/ddab215.

